# Activity and cytosolic Na^+^ regulates synaptic vesicle endocytosis

**DOI:** 10.1101/2019.12.27.889790

**Authors:** Yun Zhu, Dainan Li, Hai Huang

## Abstract

Retrieval of synaptic vesicles via endocytosis is essential for maintaining sustained synaptic transmission, especially for neurons that fire action potentials at high frequencies. However, how activity regulates synaptic vesicles recycling is largely unknown. Here we report that Na^+^ substantially accumulated in the mouse calyx of Held terminals during repetitive high-frequency spiking. Elevated presynaptic Na^+^ accelerated both slow and rapid forms of endocytosis and facilitated endocytosis overshoot but did not affect the readily releasable pool size, Ca^2+^ influx, or exocytosis. To examine whether this facilitation of endocytosis is related to the Na^+^-dependent vesicular content change, we dialyzed increasing concentrations of glutamate into the presynaptic cytosol or blocked the vesicular glutamate uptake with bafilomycin and found the rate of endocytosis was not affected by regulating the glutamate content in the presynaptic terminal. Endocytosis is critically dependent on intracellular Ca^2+^, and the activity of Na^+^/Ca^2+^ exchanger (NCX) may be altered when the Na^+^ gradient is changed. However, neither NCX blocker nor change of extracellular Na^+^ concentration affected the endocytosis rate. Moreover, two-photon Ca^2+^ imaging showed that presynaptic Na^+^ did not affect the action potential-evoked intracellular Ca^2+^ transient and decay. Therefore, we revealed a novel mechanism of cytosolic Na^+^ in accelerating vesicle endocytosis. During high-frequency synaptic transmission, when large amounts of synaptic vesicles are fused, Na^+^ accumulated in terminals, facilitated vesicle recycling and sustained reliable synaptic transmission.

## INTRODUCTION

At chemical synapses, the fusion of synaptic vesicles with the presynaptic plasma membrane releases vesicular neurotransmitter contents for exerting various functions. Following exocytosis, membrane lipid bilayer and proteins are retrieved through endocytosis to form new vesicles (Sudhof, 2004; Kononenko and Haucke, 2015; Soykan et al., 2017). Different modes of endocytosis have been observed in central synapses, including the classical clathrin-dependent endocytosis, kiss-and-run, and bulk exocytosis (Hosoi et al., 2009; Wu et al., 2009; Yamashita et al., 2010). Endocytosis in different conditions differs substantially in speed and amount. Slow endocytosis is mediated by a classical, clathrin- and dynamin-dependent endocytosis with a decay time constant (τ) of ∼10-30 s, serving as the predominant mode for synaptic vesicle recycling during low-intensity activity (Granseth et al., 2006). The clathrin-independent, dynamin-dependent rapid endocytosis, with a τ within a few seconds, has been assumed to reflect kiss-and-run that involves rapid fusion pore opening and closure of the same vesicles after stronger stimulation (Ales et al., 1999; Wu et al., 2005). Bulk endocytosis occurs when large endosome-like structures are internalized from presynaptic plasma membrane during high intensity firing activity (Holt et al., 2003; Wu and Wu, 2007). Moreover, during endocytosis overshoot, endocytosis may retrieve more membrane than exocytosis, which may be induced with intensive stimulation and large Ca^2+^ influx (Renden and von Gersdorff, 2007; Xue et al., 2012).

Accumulating evidence indicated that synaptic vesicles exocytosis and endocytosis are tightly coupled both temporally and spatially, and their timely coupling is essential for synaptic function and structural stability, but the underlying mechanism for this coupling is under debate. Ca^2+^ represents one prevalent possibility. Vesicle exocytosis is directly triggered by Ca^2+^, while how Ca^2+^ regulates endocytosis is complex and varies greatly in response to distinct neuronal activity (Leitz and Kavalali, 2016), making it complicated to coordinate endocytosis speed in an activity-dependent manner. Here we showed that action potential firing substantially elevated presynaptic cytosolic Na^+^ in the mouse calyx of Held, a giant glutamatergic terminal in the auditory brainstem that fires continuous action potentials at high-frequency up to hundreds of Hertz, in a frequency-dependent manner. Cytosolic Na^+^ accelerated both slow and rapid forms of endocytosis and facilitated endocytosis overshoot. The Na^+^ effect on endocytosis was not through affecting Ca^2+^ influx or intracellular Ca^2+^ transients, nor through regulating vesicular glutamate contents. Therefore, when large amounts of synaptic vesicles are fused during high-frequency synaptic transmission, Na^+^ accumulated into presynaptic cytosol and facilitated vesicle recycling. Since the cytosolic Na^+^ concentration is correlated with spike intensity and durations, it may work as a signal to coordinate vesicle endocytosis and recycling according to the level of endocytosis.

## RESULTS

### Spikes activity promotes presynaptic cytosolic Na^+^ concentration

Presynaptic Na^+^ changes during action potential (AP) firing were assayed using Na^+^ imaging with two-photon laser scanning microscopy. Calyces of Held were loaded via whole-cell recordings with the Na^+^ indicator SBFI and the volume marker Alexa 594 (Fig. 1A). Standard calibration methods were used to measure the absolute [Na^+^] (Supplemental Figure 1) (Rose, 2012; Huang and Trussell, 2014). The presynaptic [Na^+^] in the resting state was 15.8 ± 2.1 mM (n = 4). APs were evoked by afferent fiber stimulation and propagated to the presynaptic terminals. Upon 100 Hz stimulation for 1 s, presynaptic cytosolic [Na^+^] increased by 5.7 ± 1.1 mM, which decayed to the control level with a time constant of 12.2 ± 1.0 s (n = 5; Fig. 1B). After ensuring reliable AP evoked by afferent fiber stimulation, the recording pipettes were subsequently detached from the calyces, and the Na^+^ signals upon stimulations at different frequencies were measured. Spiking at 10 Hz for 60 s reversibly increased the [Na^+^] by 12.2 ± 1.7 mM (n = 7; Fig. 1C). Increasing the spike frequency gradually increased the Na^+^ transients, 20 Hz for 30 s increased the [Na^+^] by 16.3 ± 2.0 mM (n = 8; Fig. 1D) and 100 Hz for 20 s increased the [Na^+^] by 55.6 ± 5.9 mM (n = 8; Fig. 1E). Therefore, spike activities efficiently increase the presynaptic cytosolic Na^+^ concentration in a frequency-dependent manner.

**Figure 1.**
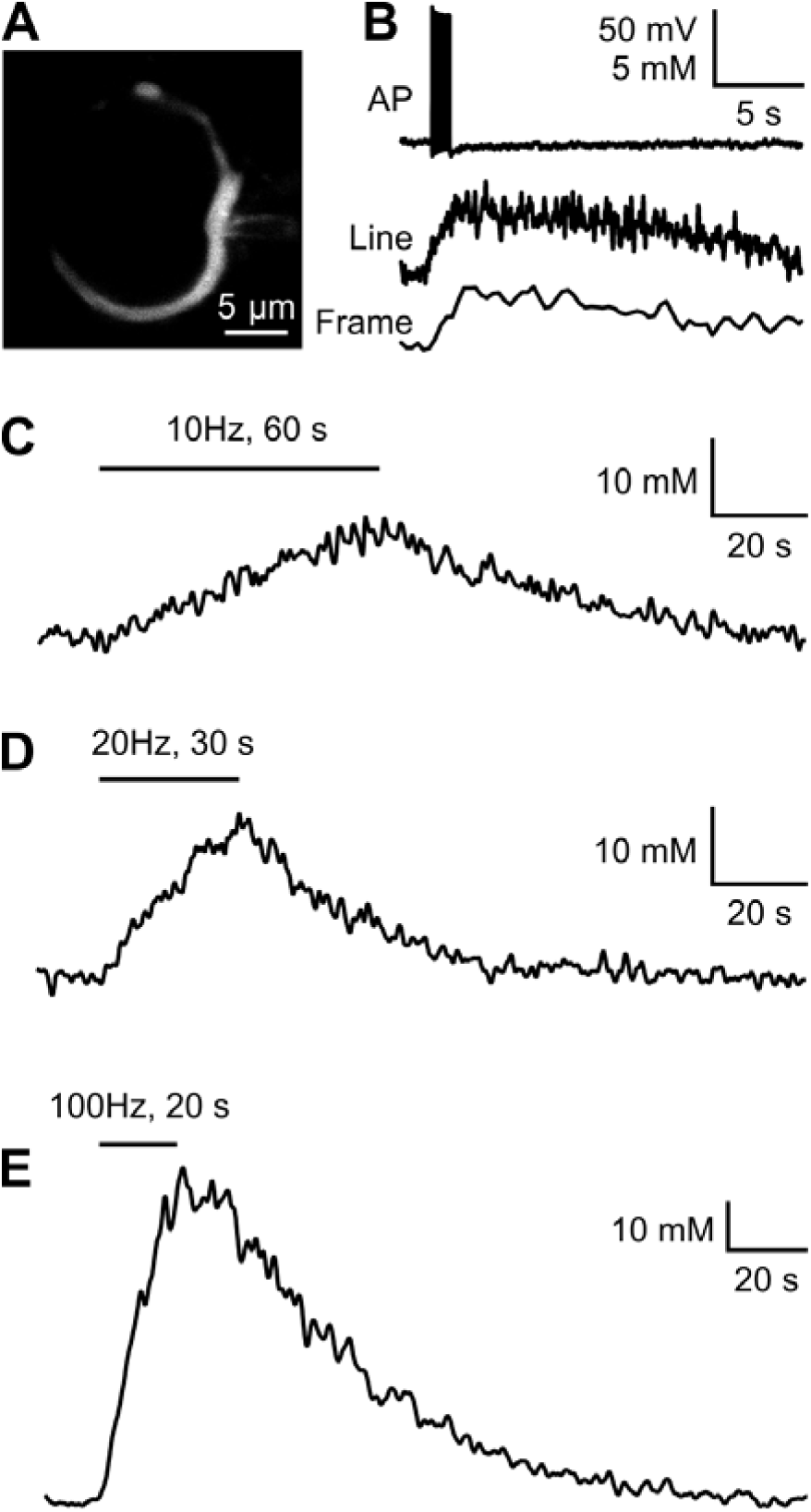
Presynaptic spikes control cytosolic Na^+^ concentration. (A) A single optical section of a calyx of Held filled with SBFI and Alexa 594 using two-photon microscopy (B) Spikes were evoked at 100 Hz for 1 s by afferent fiber stimulation. An increase in [Na^+^] was visualized with SBFI under line-scan and frame-scan modes. (C-E) After detaching recording pipettes from calyces following dialysis with SBFI and Alexa 594, Na^+^ signals in response to increasing spike frequencies were measured. Spikes at 10 Hz (C), 20 Hz (D), and 100 Hz (E) induced different changes in intracellular Na^+^ concentration.

### Presynaptic cytosolic Na^+^ facilitates slow endocytosis

Capacitance measurements were made at the calyx of Held terminals to examine whether presynaptic Na^+^ influences vesicle endocytosis and recycling. Endocytosis of the calyces depends on activity; mild stimulation triggers slow clathrin-dependent endocytosis while strong stimulation induces an additional clathrin-independent rapid form of endocytosis (Wu et al., 2005; Hosoi et al., 2009; Wu et al., 2009; Yamashita et al., 2010). We measured the whole-cell capacitance under 1.2 mM extracellular Ca^2+^ at 32°C with pipette solutions containing 0, 10 or 40 mM Na^+^. Voltage depolarizations from –80 mV to +10 mV for 40 ms (depol_40ms_) induced a Ca^2+^ influx and triggered an increase in membrane capacitance (*ΔC*_m_), followed by a slow decay of *C*_m_ toward to the prestimulus level (Fig. 2). Because *C*_m_ is proportional to the membrane surface area, *ΔC*_m_ reflects exocytosis and *C*_m_ decay reflects endocytosis (Sun and Wu, 2001). This protocol is sufficient to release of readily releasable vesicles (Wu and Borst, 1999; Fedchyshyn and Wang, 2005; Renden and von Gersdorff, 2007), thus Δ*C*_m_ reflects the size of readily releasable pool. We found that the Ca^2+^ currents (P = 0.87) and Δ*C*_m_ (P = 0.47; ANOVA test) were not different in recordings with 0, 10, or 40 mM Na^+^ (Fig. 2C-D), while presynaptic Na^+^ showed profound effects on endocytosis rate (Fig. 2E). When the terminal was dialyzed with 10mM Na^+^, the *C*_m_ decay could be fitted with monoexponential functions with a time constants (τ) of 15.3 ± 0.9 s (n = 9). With the 40 mM Na^+^ solution presynaptically, the endocytosis rate accelerated, and the time constants reduced to 10.4 ± 1.0 s (n = 10, P = 0.002). Upon dialysis with Na^+^-free solution, the endocytosis became slower, resulting in a time constant of 25.1 ± 1.9 s (n = 9, P = 0.0002; unpaired t-test). Thus, presynaptic cytosolic Na^+^ accelerates slow endocytosis at mild stimulation.

**Figure 2.**
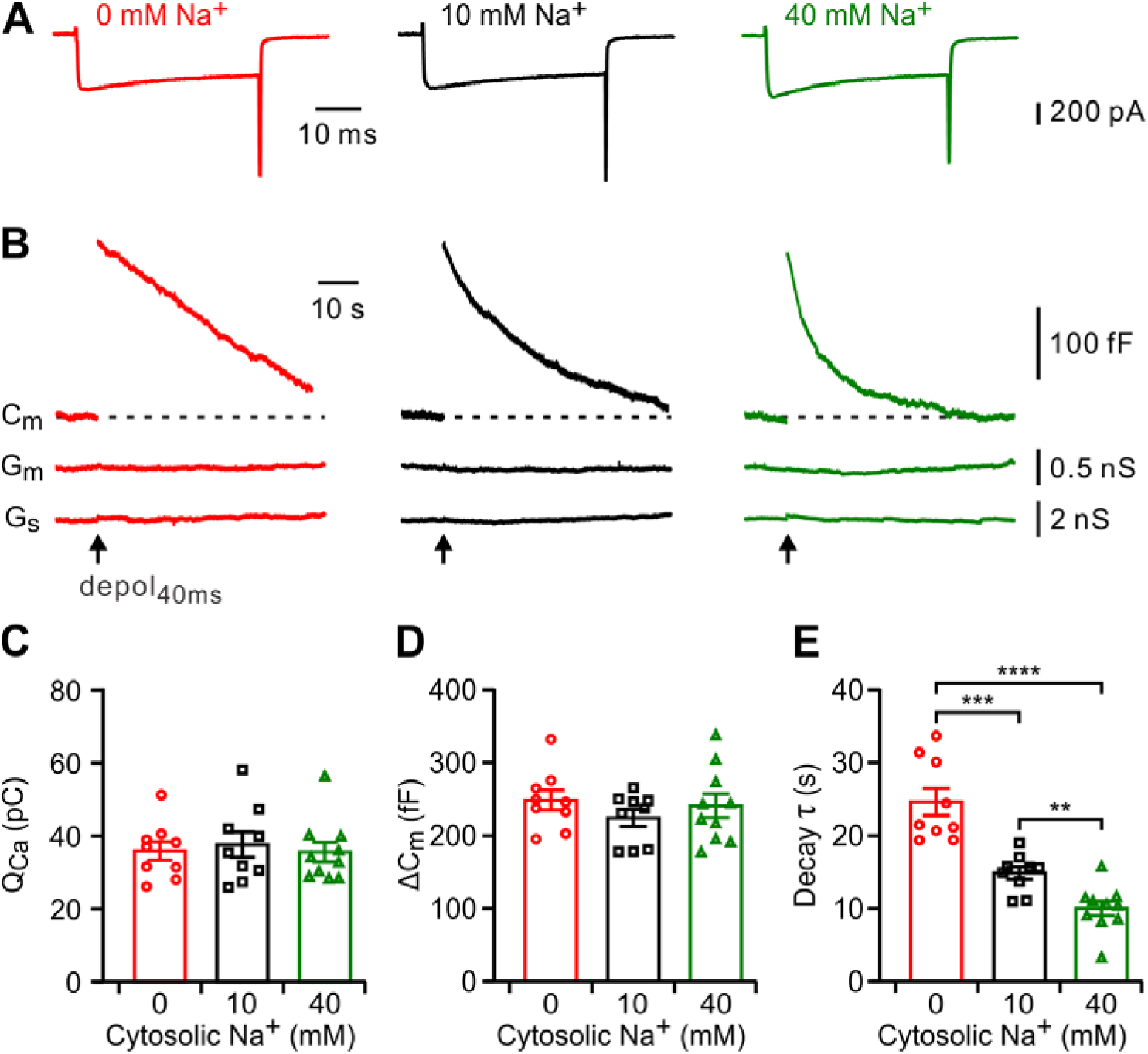
Presynaptic Na^+^ facilitates slow endocytosis without affecting exocytosis. (A) Sampled presynaptic Ca^2+^ currents induced by 40 ms depolarizations from –80 mV to +10 mV (depol_40ms_) with presynaptic pipette solutions containing 0 mM (left), 10 mM (middle), or 40 mM Na^+^ (right). (B) Corresponding membrane capacitance (C_m_) recordings showing exocytosis and endocytosis induced by depol_40ms_. Membrane conductance (G_m_) and series conductance (G_s_) are included to confirm the recording stability. (C-E) Group data show that the effects of cytosolic Na^+^ on Ca^2+^ charge (C), capacitance jump (*ΔC*_m_, D), and the time constant of capacitance decay (τ, E). **P < 0.01, ***P < 0.001, ****P < 0.0001, unpaired t-test. Error bars, ±SEM.

### Cytosolic Na^+^ facilitates rapid endocytosis

Stronger stimulation induces an additional clathrin-independent rapid form of endocytosis along with the slow endocytosis (Ales et al., 1999; Wu et al., 2005), we next investigated whether Na^+^ modulates the rapid form of endocytosis. With 10 mM [Na^+^] in the presynaptic solution, 10 pulses of 20 ms depolarization from -80 mV to +10 mV at 10 Hz (depol_20ms_×10) evoked a *ΔC*_m_ of 1002 ± 55 fF followed by a *C*_m_ decay that can be fitted with a double exponential whose fast and slow time constants were 3.3 ± 0.3 s and of 15.1 ± 1. 9 s, respectively (Fig. 3). The fast component was 40.1 ± 2.7 % of the fit (the remainder being slow component), and the weighted mean was 11.5 ± 1.1 s (n = 11). When the terminal was dialyzed with 40 mM Na^+^, both fast and weighted endocytosis were accelerated, resulting in a fast and slow components of 1.3 ± 0.2 (36.4 ± 3.4 %) and 10. 8 ± 1.1 s and a weighted time constant of 7.6 ± 0.8 s (n = 11), while with Na^+^-free solution slowed down the both fast and slow endocytosis to 4.9 ± 0.6 s (36.6 ± 4.7 %) and 22.62± 2.1 s and a weighted time constant of 18.3 ± 1.8 s (n = 10). The evoked Δ*C*_m_ was not different among groups (Fig. 3E; P = 0.67). These results revealed that presynaptic cytosolic Na^+^ facilitates both the slow and rapid modes of synaptic vesicle endocytosis.

**Figure 3.**
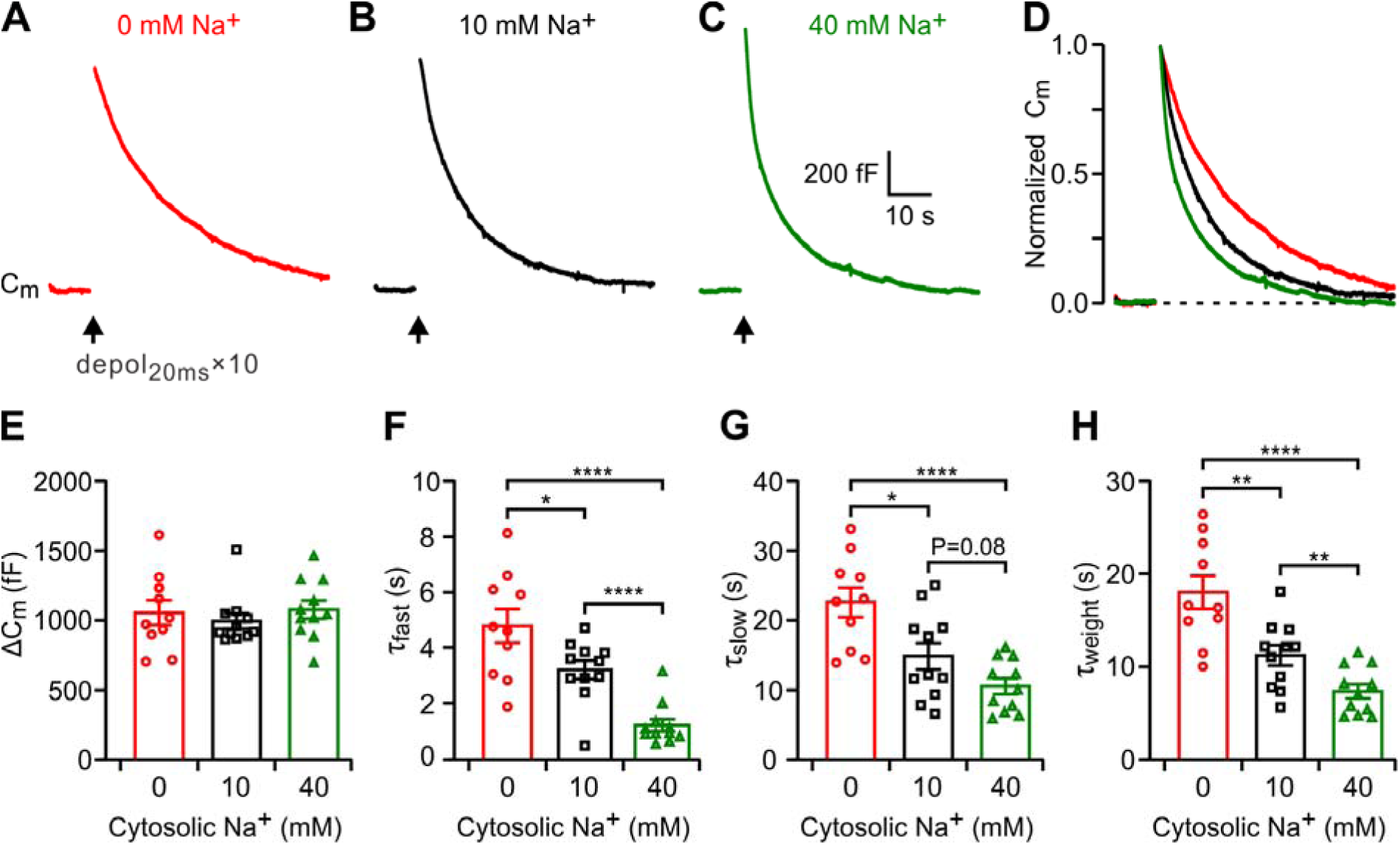
Cytosolic Na^+^ facilitates rapid endocytosis. (A-C) Sampled membrane capacitance (C_m_) recordings showing exocytosis and endocytosis induced by 10 of 20ms depolarizations (depol_20ms_×10) with pipette solutions containing 0 mM (A), 10 mM (B), or 40 mM (C) Na^+^. (D) Normalized C_m_ traces of A-C showing the Na^+^ effects on endocytosis rate. (E) Statistics for capacitance jump (*ΔC*_m_). (F-H) The capacitance decay was fitted by double exponetials. The fast (τ_fast_, F) and slow (τ_slow_, G) components, and the weighted mean (τ_weight_, H) from different cytosolic Na^+^ concentrations are compared. *P < 0.05, **P < 0.01, ****P < 0.0001, unpaired t-test. Error bars, ±SEM.

### Cytosolic Na^+^ facilitates endocytosis overshoot

During intense stimulations, such as 10 depolarization pulses of 50 ms at 10 Hz (depol_50ms_×10), the calyx terminal could retrieve more membrane area than vesicles fused. This phenomenon is defined as endocytosis overshoot and has been implicated in increasing endocytosis capacity and efficiency during high-frequency firing (Renden and von Gersdorff, 2007; Wu et al., 2009; Xue et al., 2012). We then tested whether Na^+^ influences this mode of endocytosis. Extracellular solution containing 2 mM Ca^2+^ was used in this experiment to increase the chance of endocytosis overshoot (Wu et al., 2009; Xue et al., 2012). The size of endocytosis overshoot was quantified as the capacitance value below the baseline at 40∼60 s after depolarization pulses (Wu et al., 2009). We found that increased Na^+^ concentrations facilitated the observation of endocytosis overshoot. As the [Na^+^] increased from 0 to 10, and 40 mM, the percentage of cells with overshoot increased from 28.6% (2 of 7 calyces) to 66.7% (6 of 9 calyces), and 100% (7 of 7 calyces). For the cells that showed overshoot, the overshoot ratio increased from 5.5 ± 4.4% to 14.7 ± 3.0%, and 22.8 ± 4.3% (Fig. 4; P = 0.0008, ANOVA analysis). These results indicate that [Na^+^] also facilitates endocytosis overshoot in both the percentage of cells and the amount of membrane area.

**Figure 4.**
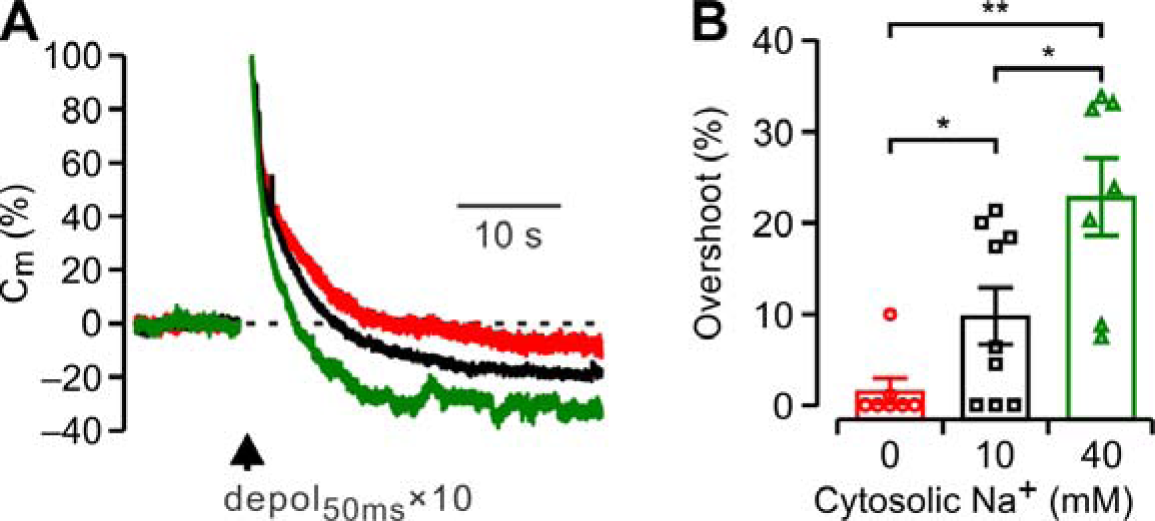
Cytosolic Na^+^ facilitates rapid endocytosis. (A) Sampled endocytosis overshoots induced by 10 of 50ms depolarizations (depol_50ms_×10) with presynaptic pipette solutions containing 0 mM (red), 10 mM (black), or 40 mM (green) Na^+^. Extracellular Ca^2+^ was elevated to 2 mM to increase the overshoot chance. (B) Statistics for endocytosis overshoot size in three groups. *P < 0.05, **P < 0.01, unpaired t-test. Error bars, ±SEM.

### Vesicular content does not affect slow or rapid endocytosis

Previous study of hippocampal cultures showed that cytosolic transmitter concentration, which rapidly controls the vesicular neurotransmitter contents (Hori and Takahashi, 2012; Apostolides and Trussell, 2013), regulates vesicle cycling at hippocampal GABAergic terminals (Wang et al., 2013). Since presynaptic Na^+^ facilitates vesicular glutamate transport and higher cytosolic [Na^+^] increases the vesicle glutamate contents (Huang and Trussell, 2014), we performed two lines of experiment to test whether the effects of Na^+^ on endocytosis is through regulating vesicular contents (Fig.5). First, we performed capacitance measurement under different concentrations of cytosolic glutamate (0, 5, or 50 mM), which allows manipulating the vesicular glutamate content (Hori and Takahashi, 2012), while keeping the Na^+^ constant at 10 mM. Slow and rapid endocytosis were evoked by the weak (depol_40ms_, Fig. 5A) and intense (depol_20ms_×10, Fig. 5E) stimulations, respectively. Our results showed that neither slow (P = 0.82, ANOVA test, Fig. 5D) nor rapid (P = 0.74, ANOVA test, Fig. 5H) mode of endocytosis was influenced by the cytosolic glutamate concentration. Second, we tested the effects of bafilomycin, a V-ATPase inhibitor that impedes vesicle acidification and neurotransmitter upload on endocytosis triggered by the same protocols (Fig. 5B,F). The results showed that 2 µM bafilomycin did not affect either slow (P = 0.30, unpaired t-test, Fig. 5D) or rapid endocytosis (P = 0.56, unpaired t-test, Fig. 5H). The *ΔC*_m_ were not different among all groups (Fig. 5C,G). The results indicates that the vesicular content had no significant effects on endocytosis or exocytosis.

**Figure 5.**
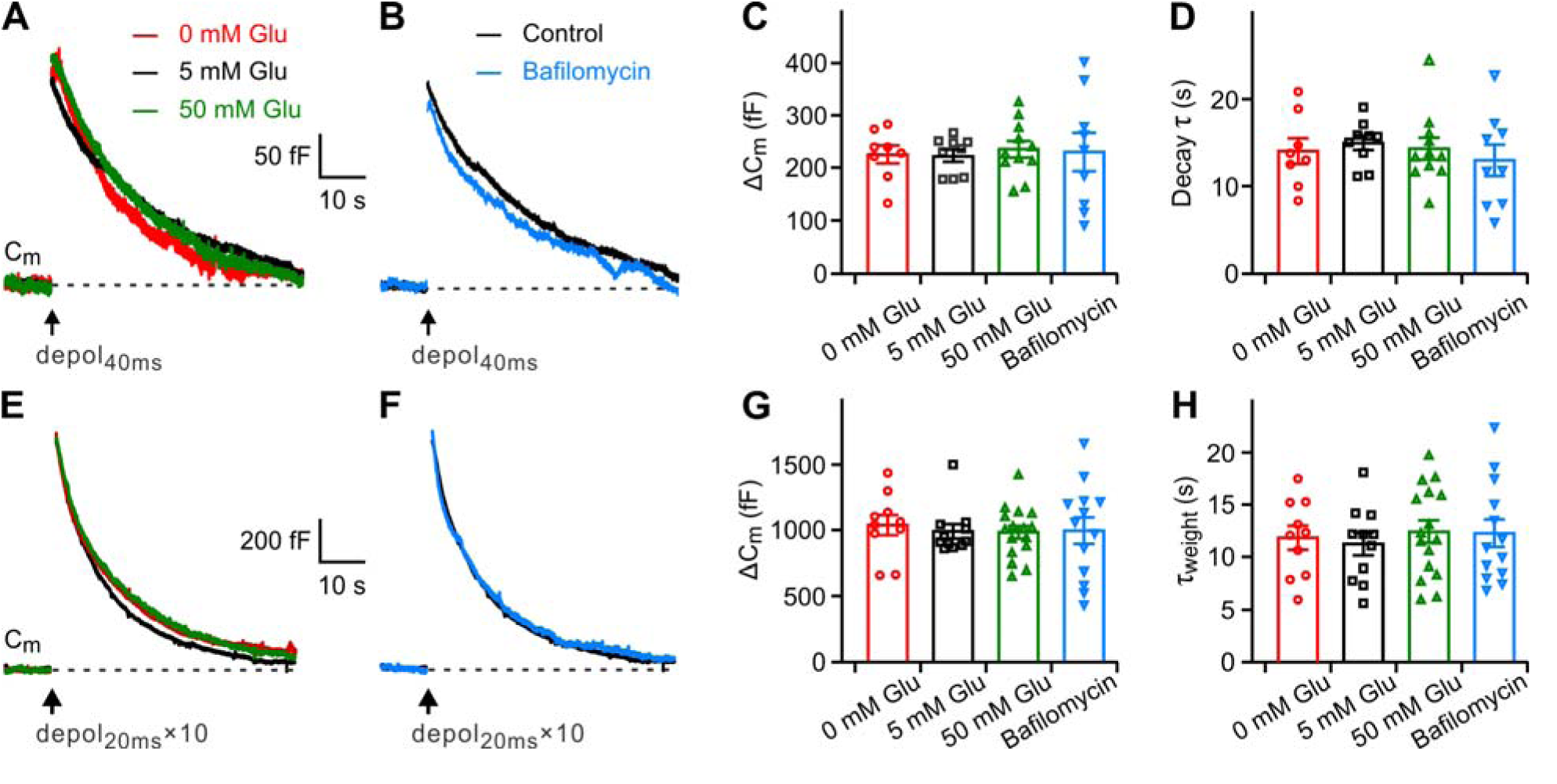
Vesicular content does not affect endocytosis. (A-B) Slow endocytosis induced by depol_40ms_ when pipette solutions contained 0, 5, or 50 mM glutamate (A), or after incubating bafilomycine A1 (B). (C-D) Neither glutamate nor bafilomycin affects the capacitance jump (*ΔC*_m_, C) or the time constant of capacitance decay τ (E). (E-H) Similar to A-D, while rapid endocytosis were induced by depol_20ms_×10. Error bars, ±SEM.

### Cytosolic Na^+^ facilitated endocytosis is independent of NCX activity

It has been shown that plasma membrane Na^+^/Ca^2+^ exchanger (NCX) is involved in maintaining presynaptic Ca^2+^ homeostasis (Kim et al., 2005; Lee and Kim, 2015), while Ca^2+^ plays specific roles in synaptic vesicle exocytosis (Leitz and Kavalali, 2016). To test whether presynaptic cytosolic Na^+^ affects the NCX function, we lowered the extracellular Na^+^ concentration to mimic the Na^+^ concentration gradient change in elevated intracellular Na^+^ conditions. As shown in Fig. 6A, the endocytosis rate was not changed with lowering extracellular Na^+^ for 50 mM (P = 0.50, n = 6). We also tested whether inhibiting NCX activity affects the endocytosis rate. As shown in Fig. 6B, incubation with the NCX inhibitor KB-R7943 (20 µM) slightly but not significantly slowed the rate of endocytosis (P = 0.14; n = 8). If Na^+^ regulates endocytosis is through slowing NCX-dependent Ca^2+^ extrusion, one would however expect an acceleration of endocytosis by NCX blocker. Therefore, the presynaptic Na^+^ regulated vesicle recycling is not mediated by modulating NCX function or Ca^2+^ extrusion.

**Figure 6.**
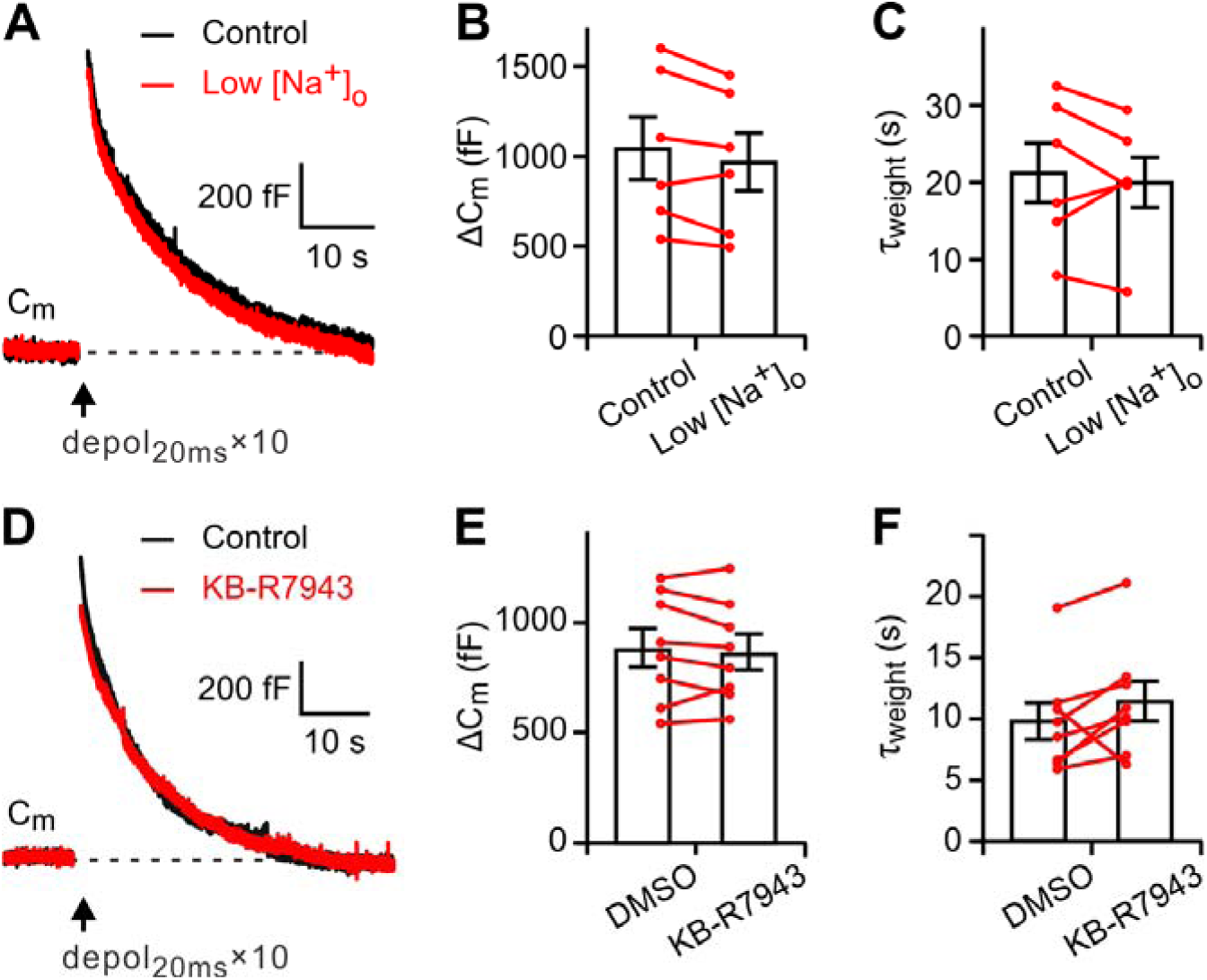
Facilitation of endocytosis by Na^+^ is not through affecting Na^+^/Ca^2+^ activity. (A, D) Sampled C_m_ recordings showing exocytosis and endocytosis induced by 10 of 20ms depolarizations (depol_20ms_×10) with in control and with low extracellular Na^+^ concentration (A) or in the presence of NCX blocker KB-R7943 (D). (B-C) Low extracellular Na^+^ does not affects the capacitance jump (*ΔC*_m_, B) or the weighted time constant of capacitance decay (τ_weight_, C). (E-F) KB-R7943 does not affects *ΔC*_m_ (E) or τ_weight_ (F). Error bars, ±SEM.

### Cytosolic Na^+^ does not affect spike-evoked intracellular Ca^2+^ rise and decay

To test whether cytosolic Na^+^ affects intracellular Ca^2+^ dynamics during spiking activity, we made two-photon Ca^2+^ imaging with Fluo-5F loaded into the presynaptic terminal (Fig. 7). Calyces were recorded with pipette solutions containing 0, 10 or 40 mM Na^+^. A burst of 10 APs evoked a rapid Ca^2+^ rise of similar concentrations at all concentrations of Na^+^ tested, as indicating by the fluorescence increases of 29.3 ± 0.9% of (G/R)_max_ in Na^+^-free(n = 8), 30.3 ± 1.5% in 10 mM Na^+^ (n = 7), and 31.3 ± 1.6% in 40 mM Na^+^ (n = 6) pipette solutions (Fig. 7C). The Ca^2+^ signals decayed to the background level within seconds and were not different among groups. The Ca^2+^ decay time constants were 0.56 ± 0.06 s for 0 mM Na^+^, 0.60 ± 0.07 s for 10 mM Na^+^ and 0.61 ± 0.05 s for 40 mM Na^+^ solutions (P = 0.80, ANOVA test; Fig. 7D). Thus, the mechanism of presynaptic Na^+^ facilitating endocytosis is unlikely through Ca^2+^-dependent pathway.

**Figure 7.**
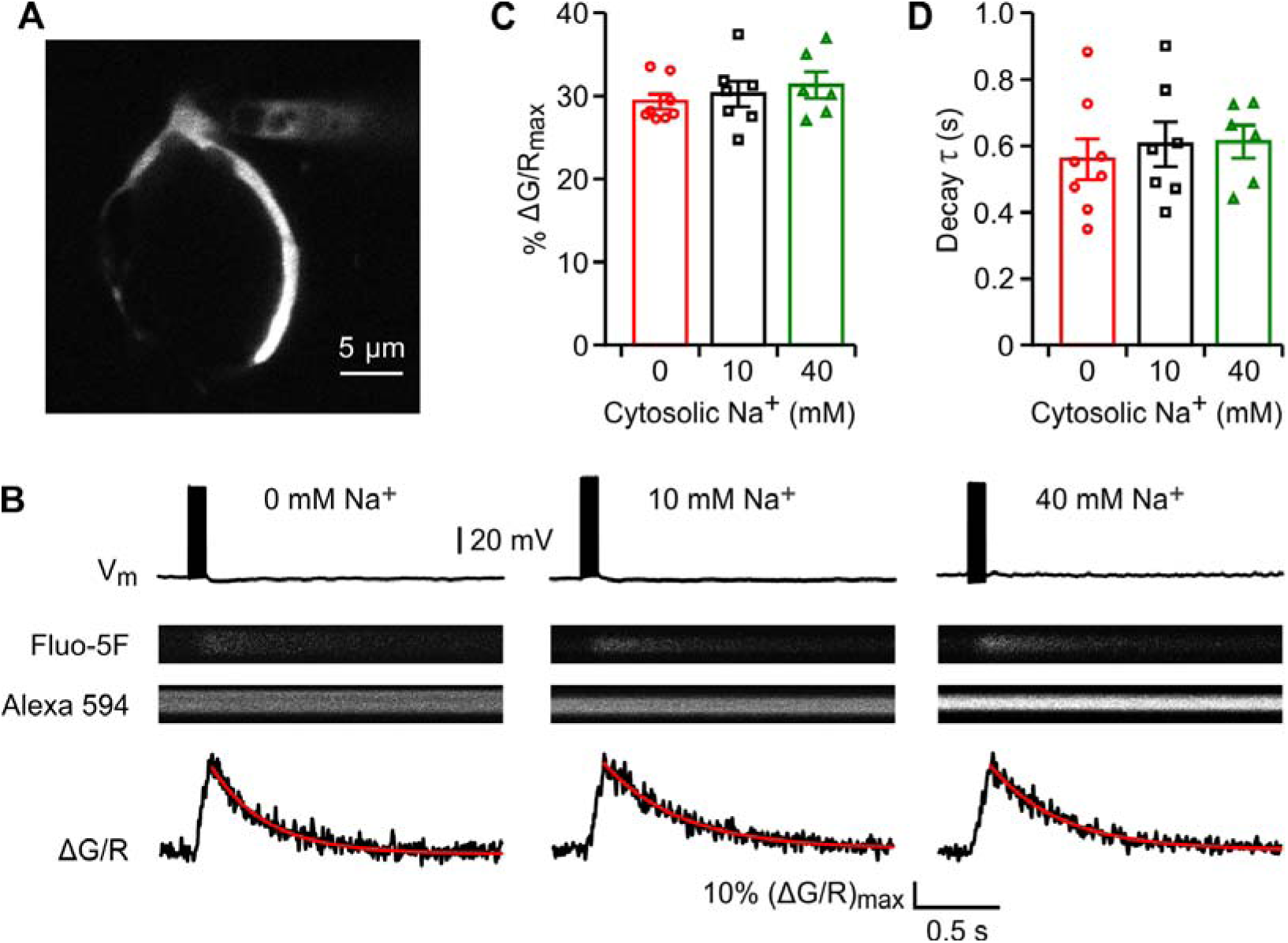
Cytosolic Na^+^ does not affect spike-evoked Ca^2+^ transients and decay. (A) Single optic section of the calyx with attached patch pipette. (B) Presynaptic Ca^2+^ transients induced by 10 spikes at 100 Hz when dialyzed with pipette solution containing 0mM, 10 mM, or 40 mM Na^+^. (C-D) Summary plots of relative Ca^2+^ rise and decay time with different intracellular Na^+^ concentrations. Error bars, ±SEM.

## DISCUSSION

Here we showed that intracellular Na^+^ concentration at the presynaptic terminal increased rapidly during spike activity (Fig. 1). This increased [Na^+^] accelerated slow (Fig. 2) and rapid endocytosis (Fig.3), as well as facilitated endocytosis overshoot (Fig. 4). The modulation of endocytosis by Na^+^ was unlikely related to vesicular glutamate contents (Fig. 5), Ca^2+^ influx through NCX (Fig. 6) or intracellular Ca^2+^ transients in response to stimulation (Fig. 7). Therefore, we revealed an unappreciated role of cytosolic Na^+^ in regulation of synaptic vesicle endocytosis and recycling. When large amounts of synaptic vesicles are fused during high-frequency synaptic transmission, accumulated presynaptic Na^+^ accelerates vesicle recycling and sustains synaptic transmission, representing a novel cellular mechanism that supports reliable synaptic transmission at high-frequency in the central nervous system.

Accumulating evidence indicated that synaptic vesicles exocytosis and endocytosis are tightly coupled both temporally and spatially, and their coupling is essential for synaptic function and structural stability. During prolonged high-frequency synaptic transmission when large amounts of synaptic vesicles are fused with presynaptic membrane, endocytosis needs to be accelerated in order to replenish the vesicle pool and maintain the strength of synaptic transmission. Slow endocytosis and vesicle recycling leads to depletion of the release-ready vesicles, which has been shown as one major contribution for short-term depression (von Gersdorff and Borst, 2002; Fernandez-Alfonso and Ryan, 2004; Regehr, 2012). Different mechanisms have been observed to couple endocytosis with exocytosis, including Ca^2+^ (Sankaranarayanan and Ryan, 2001; Hosoi et al., 2009; Wu et al., 2009), membrane lipid such as PIP2 (Koch and Holt, 2012), and cytoskeleton proteins (Yuan et al., 2015; Orlando et al., 2019). Ca^2+^-dependent regulation of exo-endocytosis coupling has been prevalently studied. Exocytosis is directly triggered by Ca^2+^ and Ca^2+^-dependent regulation of endocytosis has been extensively studied (Wu et al., 2014). Ca^2+^ speeds up slow, rapid, and bulk endocytosis while buffering intracellular Ca^2+^ with chelators or reducing Ca^2+^ influx slows down endocytosis on endocrine cells and various synapses, indicating that Ca^2+^ triggers endocytosis (Sankaranarayanan and Ryan, 2001; Hosoi et al., 2009; Wu et al., 2009). However, increased Ca^2+^ influx during prolonged stimulation train slowed down endocytosis in many preparations, including chromaffin cells, neuromuscular junction, retinal ribbon terminals, and central synapses (von Gersdorff and Matthews, 1994; Wu and Betz, 1996; Sun et al., 2002; Elhamdani et al., 2006; Balaji et al., 2008; Sun et al., 2010). A recent study showed that the dynamics of Ca^2+^ concentration changes can differentially modulate endocytosis. Specifically, transient, large calcium increases trigger endocytosis while prolonged, small, global Ca^2+^ increases inhibit slow endocytosis (Wu and Wu, 2014). Therefore, regulation of endocytosis by Ca^2+^ concentration is complex and varies greatly in response to distinct neuronal activity, making it complicated to coordinate endocytosis speed in an activity-dependent manner.

Neural activity is provided by ion fluxes through voltage and neurotransmitter-gated channels, and spiking activity alters ion composition in the cytosol. Although intracellular K^+^ concentration is high and relatively stable, the Ca^2+^ and Na^+^ levels are dynamically regulated during activity. We showed that the presynaptic cytosolic Na^+^ of the mouse calyx of Held is 15.8 ± 2.1 mM, very similar to that of the rat calyx (Huang and Trussell, 2014). We previously reported that Calyx terminals express Na^+^-permeable HCN channels, and activation of HCN channels contributes to the resting Na^+^ concentration of 4.9 ± 0.5 mM (Huang and Trussell, 2014). Here we found that spikes are much more potent in regulating presynaptic Na^+^ accumulation than HCN channels. Upon 1 s of 100 Hz firing, the presynaptic [Na^+^] increased by 5.7 ± 1.1 mM (Figure 1B), which is larger than the overall contribution of voltage-gated Na^+^ channels and HCN channels at resting potential (Huang and Trussell, 2014). Increasing the spike frequency increased the Na^+^ transients proportionately. Spikes at 10 Hz for 60 s increased the [Na^+^] by 12.2 ± 1.7 mM while 30 s of spikes at 20 Hz increased the [Na^+^] by 16.3 ± 2.0 mM (Fig. 2). Since the total spike number was the same in these two experiments, the difference in [Na^+^] increase likely reflects the Na^+^ extrusion, presumably by Na^+^/K^+^-ATPase activity. The calyx recorded in brain slices does not fire spontaneously, however, it fires *in vivo* at frequencies of 71 ± 11 Hz in the absence of sound and up to 352 ± 34 Hz with 80 dB tones (Lorteije et al., 2009). The presynaptic [Na^+^] thus would be substantially higher *in vivo* than that of the slice preparations, suggesting a mechanism to strongly modulate vesicle endocytosis *in vivo*. The presynaptic cytosolic [Na^+^] is correlated well with the firing frequency and duration (Fig. 1) and thus be able to precisely reflect the vesicle release level. Previous studies of showed that cytosolic neurotransmitter concentration regulates vesicle cycling at GABAergic terminals in hippocampal cultures (Wang et al., 2013) and transmitter concentration rapidly controls the vesicular neurotransmitter contents (Hori and Takahashi, 2012; Apostolides and Trussell, 2013). A recent study showed that a Na^+^/H^+^ exchanger expressed on synaptic vesicles promotes vesicle filling with glutamate (Goh et al., 2011). Na^+^ influx through presynaptic plasma membrane HCN channels affects presynaptic Na^+^ concentration, regulates glutamate uptake, and thus controls miniature excitatory postsynaptic currents (Huang and Trussell, 2014), suggesting that Na^+^ affects endocytosis rate through changing vesicular contents. However, our results showed that neither slow nor rapid modes of endocytosis were influenced by the cytosolic glutamate concentration (Fig. 5). These results are consistent with previous experiments in the calyx of Held (Hori and Takahashi, 2012; Takami et al., 2017). Another possible mechanism for Na^+^ effects on endocytosis is through effects on [Ca^2+^]. Na^+^ has been shown to affect presynaptic Ca^2+^ homeostasis through controlling the Na^+^/Ca^2+^ exchanger activity (Kim et al., 2005; Lee and Kim, 2015), while Ca^2+^ plays specific roles in synaptic vesicle exocytosis (Leitz and Kavalali, 2016). Our results showed that changes in extracellular Na^+^ or blocking NCX activity did not affect the endocytosis rate (Fig. 6) and cytosolic Na^+^ did not affect spike-evoked Ca^2+^ transients and decay, indicating the effects of Na^+^ on vesicle endocytosis shown here are not through Ca^2+^ and Ca^2+^-dependent pathways. The underlying mechanisms need to be studied, while recent studies showed that sodium ion is involved for the activation of endocytic protein dynamin (Chappie et al., 2010) and sodium ions allosterically modulated different G-proteins (Vickery et al., 2018), suggesting the possible interaction of Na^+^ with endocytic machinery.

We showed here that presynaptic cytosolic Na^+^ accelerated both slow and rapid forms of endocytosis and facilitated endocytosis overshoot. At high-frequency synaptic transmission, large amounts of synaptic vesicles are fused and Na^+^ accumulated into presynaptic cytosol. Na^+^ facilitates vesicle endocytosis (Fig. 1-3) and vesicular glutamate upload (Huang and Trussell, 2014), thus Na^+^ works as a signal to coordinate vesicle endocytosis and vesicular glutamate upload according to the level of endocytosis, representing a novel mechanism of vesicle exo-endocytosis coupling.

## METHODS

### Slice preparation

The care and handling of animals were approved by the Institutional Animal Care and Use Committee of Tulane University and complied with U.S. Public Health Service guidelines. Coronal brainstem slices containing the medial nucleus of the trapezoid body (MNTB) were prepared from postnatal day 8-12 C57BL/6J mice of either sex similar to previously described (Zhang and Huang, 2017). Briefly, the slices were cut using a Vibratome (VT1200S, Leica) in ice-cold, low-Ca^2+^, low-Na^+^ saline contained (in mM) 230 sucrose, 10-25 glucose, 2.5 KCl, 0.1 CaCl_2_, 3 MgCl_2_, 1.25 NaH_2_PO_4_, 25 NaHCO_3_, 0.4 ascorbic acid, 3 myo-inositol, and 2 Na-pyruvate, bubbled with 95% O_2_/5% CO_2_. Slice were incubated at 32°C for 20–30 min and thereafter stored at room temperature in normal artificial cerebrospinal fluid (aCSF) contained (in mM) 125 NaCl, 10-25 glucose, 2.5 KCl, 1.2 CaCl_2_, 1.8 MgCl_2_, 1.25 NaH_2_PO_4_, 25 NaHCO_3_, 0.4 ascorbic acid, 3 *myo-*inositol, and 2 Na-pyruvate, pH 7.4, bubbled with 5% CO_2_/95% O_2_ before use.

### Membrane capacitance measurement

Slices were transferred to a recording chamber and perfused with bubbled aCSF (2-3 ml/min) warmed to ∼32°C by an in-line heater (Warner Instruments). Whole-cell recordings were made from calyces of Held with an EPC-10 USB patch-clamp amplifier with lock-in system and PatchMaster software (HEKA, Germany). For whole-cell membrane capacitance (C_m_) measurements, the sinusoidal stimulus frequency was 1 kHz and the peak-to-peak voltage was 60 mV (Lindau and Neher, 1988). To isolate presynaptic Ca^2+^ currents in voltage-clamp experiments, TEA-Cl (10 mM), 4-aminopyridine (0.5 mM) and tetrodotoxin (1 µM) were added to aCSF, substituting for NaCl with equal osmolarity. For recordings of the endocytosis overshoot, MgCl_2_ and CaCl_2_ were adjusted to 1.0 and 2.0 mM, respectively, to facilitate overshoot (Wu et al., 2009; Xue et al., 2012). To test endocytosis under lowered the extracellular Na^+^ concentration, 50 mM NaCl was replaced by LiCl with equal osmolarity (Kim et al., 2005). The pipette solution contained (in mM): 70 Cs-methanesulfonate, 20 CsCl, 10 HEPES, 0.5 EGTA, 4 Mg-ATP, 0.3 Tris_3_-GTP, 10 Tris_2_-phosphocreatine, 5 Glutamate, as well as 40 NMDG-methanesulfonate (for Na^+^-free solution), 10 Na-methanesulfonate + 30 NMDG-methanesulfonate (for 10 mM Na^+^ solution), or 40 Na-methanesulfonate (for 40 mM Na^+^ solution). For presynaptic glutamate dialysis experiments, glutamate was added to substitute methanesulfonate with equal osmolarity. Bafilomycin A1 and KB-R7943 were prepared in stock solution with DMSO and added to recording solution immediately before experiments. All solutions were adjusted to pH 7.3 with CsOH (310-315 mOsm). Tips of patch pipettes (3–5 MΩ) were coated with dental wax to reduce stray capacitance. Series resistance (< 20 MΩ) was compensated up to 60% (10 μs lag). Data were obtained 4–20 min after break-in at sampling rate of 100 kHz and filtered with an online Bessel filter at 2.9 kHz. Liquid junction potentials (10-11mV) were measured and adjusted appropriately.

### Two-photon Na^+^ and Ca^2+^-imaging

A Galvo multiphoton microscopy system (Scientifica) with a Ti:sapphire pulsed laser (Chameleon Ultra II; Coherent) was used for two-photon Na^+^ and Ca^2+^ imaging. The laser was tuned to 800 nm for Na^+^ imaging and 810 nm for Ca^2+^ imaging, and epifluorescence signals were captured through 60×, 1.0 NA objectives and a 1.4 NA oil immersion condenser (Olympus). Fluorescence was split into red and green channels using dichroic mirrors and band-pass filters. Data were collected in frame-scan or line-scan modes using SciScan (Scientifica) or ScanImage.

For Ca^2+^ imaging under different Na^+^ concentrations, pipette solution contained (in mM): 70 K-methanesulfonate, 20 KCl, 10 HEPES, 4 Mg-ATP, 0.3 Tris_3_-GTP, 10 Tris_2_-phosphocreatine, 5 Glutamate, as well as 40 NMDG-methanesulfonate (for Na^+^-free solution), 10 Na-methanesulfonate + 30 NMDG-methanesulfonate (for 10 mM Na^+^ solution), or 40 Na-methanesulfonate (for 40 mM Na^+^ solution). 250 μM Fluo-5F and 20 μM Alexa 594 were also added to the pipette solution before the experiments. Data are expressed as Δ(G/R)/(G/R)_max_ × 100%, where (G/R)_max_ was the maximal fluorescence in saturating Ca^2+^ (Spratt et al., 2019).

For Na^+^ imaging, pipette solution contained (in mM): 110 K-methanesulfonate, 20 KCl, 10 HEPES, 0.5 EGTA, 4 Mg-ATP, 0.3 Tris_3_-GTP, 5 Na_2_-phosphocreatine (290 mOsm; pH 7.3 with KOH), while 1 mM SBFI and 15 μM Alexa 594 were added. All solutions were adjusted to pH 7.3 with KOH (290 mOsm). Presynaptic spikes were evoked by afferent fiber stimulation and recorded under current-clamp, and corresponding Na^+^ and Ca^2+^ signals were recorded under two-photon imaging. To avoid the perfusion of cytoplasmic Na^+^ during resting and spike-evoked Na^+^ signals recordings, pipettes were subsequently detached after whole-cell dialysis with SBFI and Alexa 594 dyes when the fluorescence intensities are stable. After waiting for 10 min, presynaptic spikes were evoked by afferent fiber stimulation and the Na^+^ signals were measured during the process. Standard calibration methods were used to measure the absolute Na^+^ concentrations (Rose, 2012; Huang and Trussell, 2014). The solutions for *in situ* calibration of SBFI fluorescence contained (in mM): 20 KCl, 25 glucose, 10 HEPES, 130 (K-gluconate + Na-gluconate), and adjusted to pH 7.4 with KOH. 3 μM gramicidin D, 10 μM monensin, and 50 μM ouabain were added into the calibration solutions before experiments.

### Analysis

Data were analyzed using PatchMaster (HEKA), Clampfit (Molecular Devices), Igor (WaveMetrics), and Image J(NIH). The standard monoexponential functions were used to describe slow endocytosis: *f*(t) = A × e^-t/τ^ + C, where A is the capacitance amplitudes (Δ*C*_m_), and τ is the time constant. Double exponentials functions were used to describe endocytosis under strong stimulations: *f*(t) = A_1_ × e^-t/τ1^ + A_2_ × e^-t/τ2^ + C, where A_1_ and A_2_ are the amplitudes of fast and slow exponential components; and τ_1_ and τ_2_ are the time constants of each components, respectively. The weighted time constant is expressed: τ_weight_ = (A_1_ × τ_1_ + A_2_ × τ_2_) / (A_1_ + A_2_). Data were presented as the mean ± SEM. Statistical significance was established using paired and unpaired t-tests, as well as one-way ANOVA followed by Dunnett’s post hoc test as indicated, with P < 0.05 indicating a significant difference.

### Drugs

Drugs were obtained from Alomone (tetrodotoxin), LC labs (bafilomycin A1), Invitrogen (SBFI, fluo-5F, and Alexa 594), and all others from Sigma-Aldrich.

## ACKNOWLEDGEMENTS

We thank Dr. Laura Schrader and Youad Darwish for critical reading of the manuscript. Financial supported was provided by U.S. National Institutes of Health grant R01DC016324 to H. Huang.

## AUTHOR CONTRIBUTIONS

Y.Z., D.L. and H.H. designed, performed and analyzed experiments. Y.Z. and H.H. wrote the manuscript. H.H. conceived and supervised the research project.

---

**Supplemental Figure 1.**
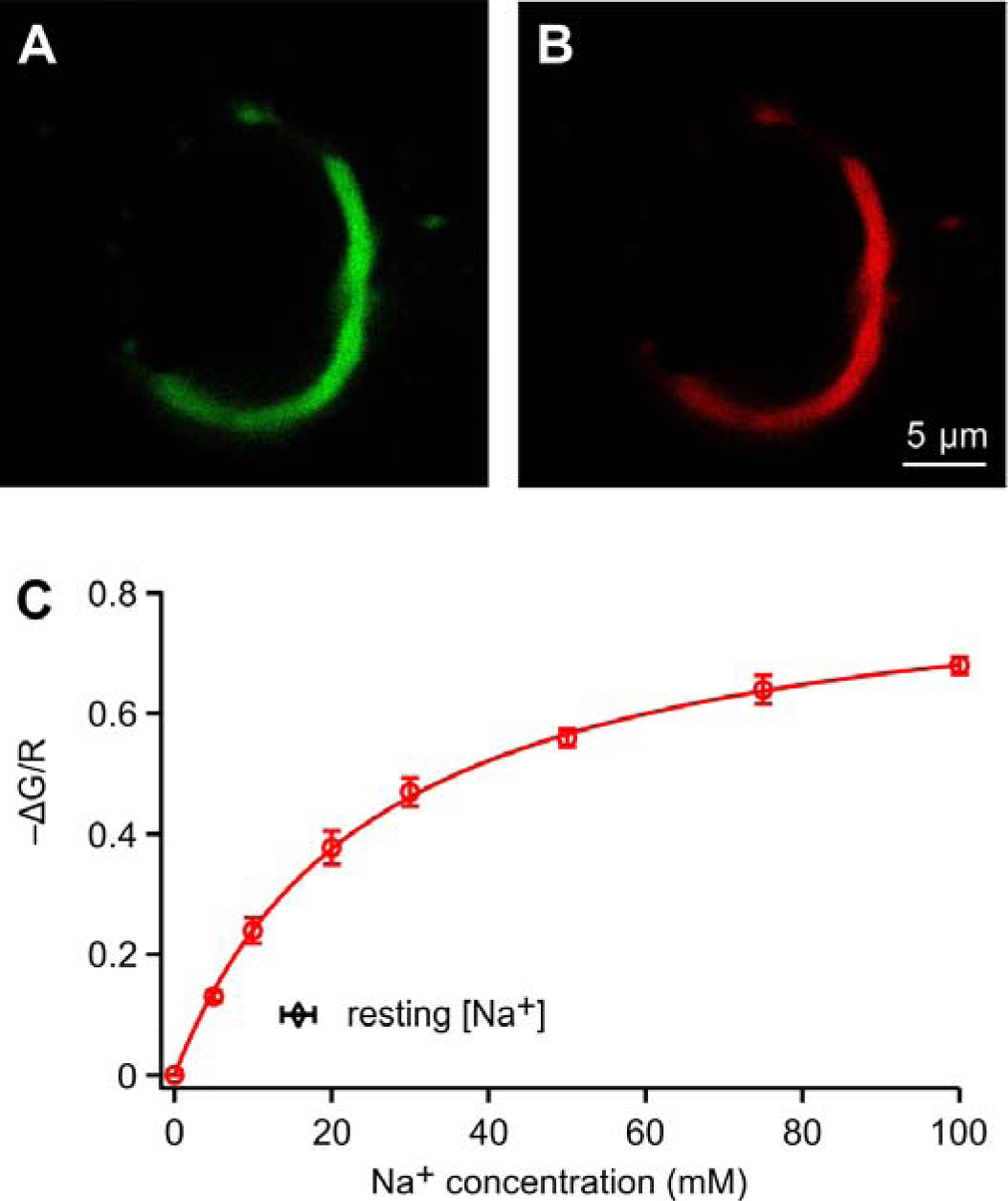
Calibration of SBFI fluorescence with changes in [Na^+^]_i_. (A-B) A single optical section of fluorescence from SBFI (A) and Alexa 594 (B) in a calyx of Held terminal. Scale bars are 5 µm for both panels. (C) Change in fluorescence with [Na^+^]_i_ (n = 5). The incubation solutions containing ionophores and ouabain (see Experimental Procedures / Materials and Methods), changes in the extracellular [Na^+^] from 0 to 100 mM evoked decreases in the SBFI fluorescence. Data are normalized and fitted by the equation: (ΔG/R) = (G/R)_max_ × [Na^+^]_i_ / ([Na^+^]_i_ + K_app_), where G/R is the ratio of green fluorescence relative to red fluorescence; (ΔG/R) is the change in fluorescence ratio measured at a given [Na^+^]_i_ divided by that at 0 mM [Na^+^]_i_; (G/R)_max_ is the maximal change in fluorescence ratio and K_app_ is the apparent K_d_ of SBFI. The fitted curve yielded a K_app_ of 25.7 mM and (G/R)_max_ of 0.86 for SBFI. The resting [Na^+^]_i_ was measured and shown in the insert. Error bars, ± S.E.M.

## REFERENCE

Ales E, Tabares L, Poyato JM, Valero V, Lindau M, Alvarez de Toledo G (1999) High calcium concentrations shift the mode of exocytosis to the kiss-and-run mechanism. Nat Cell Biol 1:40–44.

Apostolides PF, Trussell LO (2013) Rapid, activity-independent turnover of vesicular transmitter content at a mixed glycine/GABA synapse. J Neurosci 33:4768–4781.

Balaji J, Armbruster M, Ryan TA (2008) Calcium control of endocytic capacity at a CNS synapse. J Neurosci 28:6742–6749.

Chappie JS, Acharya S, Leonard M, Schmid SL, Dyda F (2010) G domain dimerization controls dynamin’s assembly-stimulated GTPase activity. Nature 465:435–440.

Elhamdani A, Azizi F, Artalejo CR (2006) Double patch clamp reveals that transient fusion (kiss-and-run) is a major mechanism of secretion in calf adrenal chromaffin cells: high calcium shifts the mechanism from kiss-and-run to complete fusion. J Neurosci 26:3030–3036.

Fedchyshyn MJ, Wang LY (2005) Developmental transformation of the release modality at the calyx of Held synapse. J Neurosci 25:4131–4140.

Fernandez-Alfonso T, Ryan TA (2004) The kinetics of synaptic vesicle pool depletion at CNS synaptic terminals. Neuron 41:943–953.

Goh GY, Huang H, Ullman J, Borre L, Hnasko TS, Trussell LO, Edwards RH (2011) Presynaptic regulation of quantal size: K+/H+ exchange stimulates vesicular glutamate transport. Nat Neurosci 14:1285–1292.

Granseth B, Odermatt B, Royle SJ, Lagnado L (2006) Clathrin-mediated endocytosis is the dominant mechanism of vesicle retrieval at hippocampal synapses. Neuron 51:773–786.

Holt M, Cooke A, Wu MM, Lagnado L (2003) Bulk membrane retrieval in the synaptic terminal of retinal bipolar cells. J Neurosci 23:1329–1339.

Hori T, Takahashi T (2012) Kinetics of synaptic vesicle refilling with neurotransmitter glutamate. Neuron 76:511–517.

Hosoi N, Holt M, Sakaba T (2009) Calcium dependence of exo- and endocytotic coupling at a glutamatergic synapse. Neuron 63:216–229.

Huang H, Trussell LO (2014) Presynaptic HCN channels regulate vesicular glutamate transport. Neuron 84:340–346.

Kim MH, Korogod N, Schneggenburger R, Ho WK, Lee SH (2005) Interplay between Na+/Ca2+ exchangers and mitochondria in Ca2+ clearance at the calyx of Held. J Neurosci 25:6057–6065.

Koch M, Holt M (2012) Coupling exo- and endocytosis: an essential role for PIP(2) at the synapse. Biochim Biophys Acta 1821:1114–1132.

Kononenko NL, Haucke V (2015) Molecular mechanisms of presynaptic membrane retrieval and synaptic vesicle reformation. Neuron 85:484–496.

Lee SY, Kim JH (2015) Mechanisms underlying presynaptic Ca2+ transient and vesicular glutamate release at a CNS nerve terminal during in vitro ischaemia. J Physiol 593:2793–2806.

Leitz J, Kavalali ET (2016) Ca2+ Dependence of Synaptic Vesicle Endocytosis. Neuroscientist 22:464–476.

Lindau M, Neher E (1988) Patch-clamp techniques for time-resolved capacitance measurements in single cells. Pflugers Arch 411:137–146.

Lorteije JA, Rusu SI, Kushmerick C, Borst JG (2009) Reliability and precision of the mouse calyx of Held synapse. J Neurosci 29:13770–13784.

Orlando M, Schmitz D, Rosenmund C, Herman MA (2019) Calcium-Independent Exo-endocytosis Coupling at Small Central Synapses. Cell Rep 29:3767–3774 e3763.

Regehr WG (2012) Short-term presynaptic plasticity. Cold Spring Harb Perspect Biol 4:a005702.

Renden R, von Gersdorff H (2007) Synaptic vesicle endocytosis at a CNS nerve terminal: faster kinetics at physiological temperatures and increased endocytotic capacity during maturation. J Neurophysiol 98:3349–3359.

Rose CR (2012) Two-photon sodium imaging in dendritic spines. Cold Spring Harb Protoc 2012:1161–1165.

Sankaranarayanan S, Ryan TA (2001) Calcium accelerates endocytosis of vSNAREs at hippocampal synapses. Nat Neurosci 4:129–136.

Soykan T, Kaempf N, Sakaba T, Vollweiter D, Goerdeler F, Puchkov D, Kononenko NL, Haucke V (2017) Synaptic Vesicle Endocytosis Occurs on Multiple Timescales and Is Mediated by Formin-Dependent Actin Assembly. Neuron 93:854–866 e854.

Spratt PWE, Ben-Shalom R, Keeshen CM, Burke KJ, Jr., Clarkson RL, Sanders SJ, Bender KJ (2019) The Autism-Associated Gene Scn2a Contributes to Dendritic Excitability and Synaptic Function in the Prefrontal Cortex. Neuron 103:673–685 e675.

Sudhof TC (2004) The synaptic vesicle cycle. Annual review of neuroscience 27:509–547.

Sun JY, Wu LG (2001) Fast kinetics of exocytosis revealed by simultaneous measurements of presynaptic capacitance and postsynaptic currents at a central synapse. Neuron 30:171–182.

Sun JY, Wu XS, Wu LG (2002) Single and multiple vesicle fusion induce different rates of endocytosis at a central synapse. Nature 417:555–559.

Sun T, Wu XS, Xu J, McNeil BD, Pang ZP, Yang W, Bai L, Qadri S, Molkentin JD, Yue DT, Wu LG (2010) The role of calcium/calmodulin-activated calcineurin in rapid and slow endocytosis at central synapses. J Neurosci 30:11838–11847.

Takami C, Eguchi K, Hori T, Takahashi T (2017) Impact of vesicular glutamate leakage on synaptic transmission at the calyx of Held. J Physiol 595:1263–1271.

Vickery ON, Carvalheda CA, Zaidi SA, Pisliakov AV, Katritch V, Zachariae U (2018) Intracellular Transfer of Na(+) in an Active-State G-Protein-Coupled Receptor. Structure 26:171–180 e172.

von Gersdorff H, Matthews G (1994) Inhibition of endocytosis by elevated internal calcium in a synaptic terminal. Nature 370:652–655.

von Gersdorff H, Borst JG (2002) Short-term plasticity at the calyx of held. Nat Rev Neurosci 3:53–64.

Wang L, Tu P, Bonet L, Aubrey KR, Supplisson S (2013) Cytosolic transmitter concentration regulates vesicle cycling at hippocampal GABAergic terminals. Neuron 80:143–158.

Wu LG, Betz WJ (1996) Nerve activity but not intracellular calcium determines the time course of endocytosis at the frog neuromuscular junction. Neuron 17:769–779.

Wu LG, Borst JG (1999) The reduced release probability of releasable vesicles during recovery from short-term synaptic depression. Neuron 23:821–832.

Wu LG, Hamid E, Shin W, Chiang HC (2014) Exocytosis and endocytosis: modes, functions, and coupling mechanisms. Annu Rev Physiol 76:301–331.

Wu W, Wu LG (2007) Rapid bulk endocytosis and its kinetics of fission pore closure at a central synapse. Proc Natl Acad Sci U S A 104:10234–10239.

Wu W, Xu J, Wu XS, Wu LG (2005) Activity-dependent acceleration of endocytosis at a central synapse. J Neurosci 25:11676–11683.

Wu XS, Wu LG (2014) The yin and yang of calcium effects on synaptic vesicle endocytosis. J Neurosci 34:2652–2659.

Wu XS, McNeil BD, Xu J, Fan J, Xue L, Melicoff E, Adachi R, Bai L, Wu LG (2009) Ca(2+) and calmodulin initiate all forms of endocytosis during depolarization at a nerve terminal. Nat Neurosci 12:1003–1010.

Xue L, McNeil BD, Wu XS, Luo F, He L, Wu LG (2012) A membrane pool retrieved via endocytosis overshoot at nerve terminals: a study of its retrieval mechanism and role. J Neurosci 32:3398–3404.

Yamashita T, Eguchi K, Saitoh N, von Gersdorff H, Takahashi T (2010) Developmental shift to a mechanism of synaptic vesicle endocytosis requiring nanodomain Ca2+. Nat Neurosci 13:838–844.

Yuan T, Liu L, Zhang Y, Wei L, Zhao S, Zheng X, Huang X, Boulanger J, Gueudry C, Lu J, Xie L, Du W, Zong W, Yang L, Salamero J, Liu Y, Chen L (2015) Diacylglycerol Guides the Hopping of Clathrin-Coated Pits along Microtubules for Exo-Endocytosis Coupling. Dev Cell 35:120–130.

Zhang Y, Huang H (2017) SK Channels Regulate Resting Properties and Signaling Reliability of a Developing Fast-Spiking Neuron. J Neurosci 37:10738–10747.

